# Complex haplotypes of metabolizing *GSTM1* gene deletion harbors signatures of a selective sweep in East Asian populations

**DOI:** 10.1101/287417

**Authors:** M. Saitou, Y. Satta, O. Gokcumen

## Abstract

The deletion of the metabolizing Glutathione S-transferase Mu 1 (*GSTM1*) gene was previously associated with multiple cancers, metabolic and autoimmune disorders, as well as drug response. It is unusually common, with allele frequency reaching up to 75% in some human populations. Such high allele frequency of a derived allele with apparent impact on an otherwise conserved gene is a rare phenomenon. To investigate the evolutionary history of this locus, we analyzed 310 genomes using population genetics tools. Our analysis revealed a surprising lack of linkage disequilibrium between the deletion and the flanking single nucleotide variants in this locus, indicating gene conversion events. Tests that measure extended homozygosity and rapid change in allele frequency identified signatures of an incomplete soft-sweep in the locus. Using empirical approaches, we identified the *Tanuki* haplogroup, which carries the *GSTM1* deletion and is found in approximately 70% of East Asian chromosomes. This haplogroup has rapidly increased its frequency in East Asian populations, contributing to a high population differentiation among continental human populations. We showed that extended homozygosity and population differentiation for this haplogroup is incompatible with simulated neutral expectations in East Asian populations. In parallel, we revealed that the *Tanuki* haplogroup is significantly associated with the expression levels of other *GSTM* genes. Collectively, our results suggest that the *Tanuki* haplogroup has likely undergone a soft sweep in East Asia with multiple functional consequences. Our study provides the necessary framework for further studies to elucidate the evolutionary reasons that maintain disease-susceptibility variants in the *GSTM1* locus.

**Lay summary:** Here, we describe the evolutionary forces that shape the variation in a genomic region, which has been associated with bladder cancer, metabolic and autoimmune disorders and response to different drugs. Our results reveal a new genetic type common in Asian populations that may have important evolutionary and biomedical implications.

## INTRODUCTION

Thousands of structural variants (SVs, *i*.*e*., deletions, duplications, inversions and translocations of large segments of DNA) comprise a major part of genetic variation among humans (Redon *et al*. 2006; Sudmant *et al*. 2015b). Several studies have shown that common SVs can have important functional effects, contributing to both normal and pathogenic phenotypic variation (Zhang *et al*. 2009; Weischenfeldt *et al*. 2013). In addition, there were multiple recent studies showing adaptive phenotypes that are underlain by SVs in humans (Perry *et al*. 2007; Girirajan *et al*. 2011). However, there is still a major gap in our understanding of the functional and evolutionary impact of SVs due to three interrelated factors. First, multiple mutational mechanisms lead to SV formation, affecting the size and sequence content of the resulting variants, as well as the rate of their formation (Hastings *et al*. 2009). Second, most of the functionally relevant SVs are in complex, segmental duplication regions (Feuk *et al*. 2006; Marques-Bonet *et al*. 2009). This genomic context complicates discovery and genotyping of SVs themselves and also leads to higher than usual false negative and false positive single nucleotide variation calls in those regions. Third, the segmental duplication regions also lead to frequent non-allelic homologous recombination events, forming new SVs and leading to frequent gene conversions. Both recurrent SVs and allele swapping due to gene conversion events break the linkage disequilibrium in such loci (Usher *et al*. 2015; Boettger *et al*. 2016) Linkage disequilibrium-based imputation is essential for many evolutionary genetic analyses (Crisci *et al*. 2013) and genome-wide association studies (Visscher *et al*. 2012). As a result, the majority of SVs cannot be imputed accurately using “tag” single nucleotide polymorphisms (Wellcome Trust Case Control Consortium *et al*. 2010) and the standard, haplotype-based neutrality tests cannot be directly conducted.

To overcome these issues, recent studies have focused on resolving complex haplotype architectures harboring SVs in a locus-specific manner. For example, the reassessment of SVs in the haptoglobin locus revealed recurrent exonic deletions that are associated with blood cholesterol levels (Boettger *et al*. 2016). Similarly, the reconstruction of the haplotype-level variation in the *GYPA* and *GYPB* (Glycophorin A and Glycophorin B) locus has revealed that a specific SV leads to resistance to malaria in African populations (Leffler *et al*. 2017). To highlight some of the negative results, careful reconstruction of the haplotype-level variation in salivary *Amylase* (Usher *et al*. 2015) and the salivary *MUC7* (Mucin7) *loci* showed that previous associations in these loci with obesity and asthma, respectively, may be false. The latter study also found evidence of archaic hominin introgression in Africa affected the structural variation in this locus (Xu *et al*. 2017b). Overall, with the availability of population-level genome variation datasets, it is possible to scrutinize the evolutionary and functional impact of SVs residing in complex regions of the genome.

In a similar vein, we focus on resolving the mechanisms and evolutionary forces that lead to the common haplotypes that carry the deletion of the glutathione S-transferase mu 1 (*GSTM1*) gene in humans (**Fig. 1A**). The whole deletion of the *GSTM1* gene, which codes for a cellular detoxifying enzyme, is evolutionarily relevant as this enzyme contributes to cellular detoxification of products generated by oxidative stress, electrophilic compounds, carcinogens, environmental toxins and therapeutic drugs (McIlwain *et al*. 2006). In particular, Rothman et al. (2010) found a significant association between the *GSTM1* deletion and bladder cancer susceptibility and there are earlier reports of the deletion allele affecting response rates to chemotherapy (Hayes and Pulford 1995). Moreover, the deletion of *GSTM1* and other variants in this locus may have a broader effect on the neighboring *GSTM* gene family, members of which have similar metabolizing functions (Hayes *et al*. 2005).

**Figure 1.**
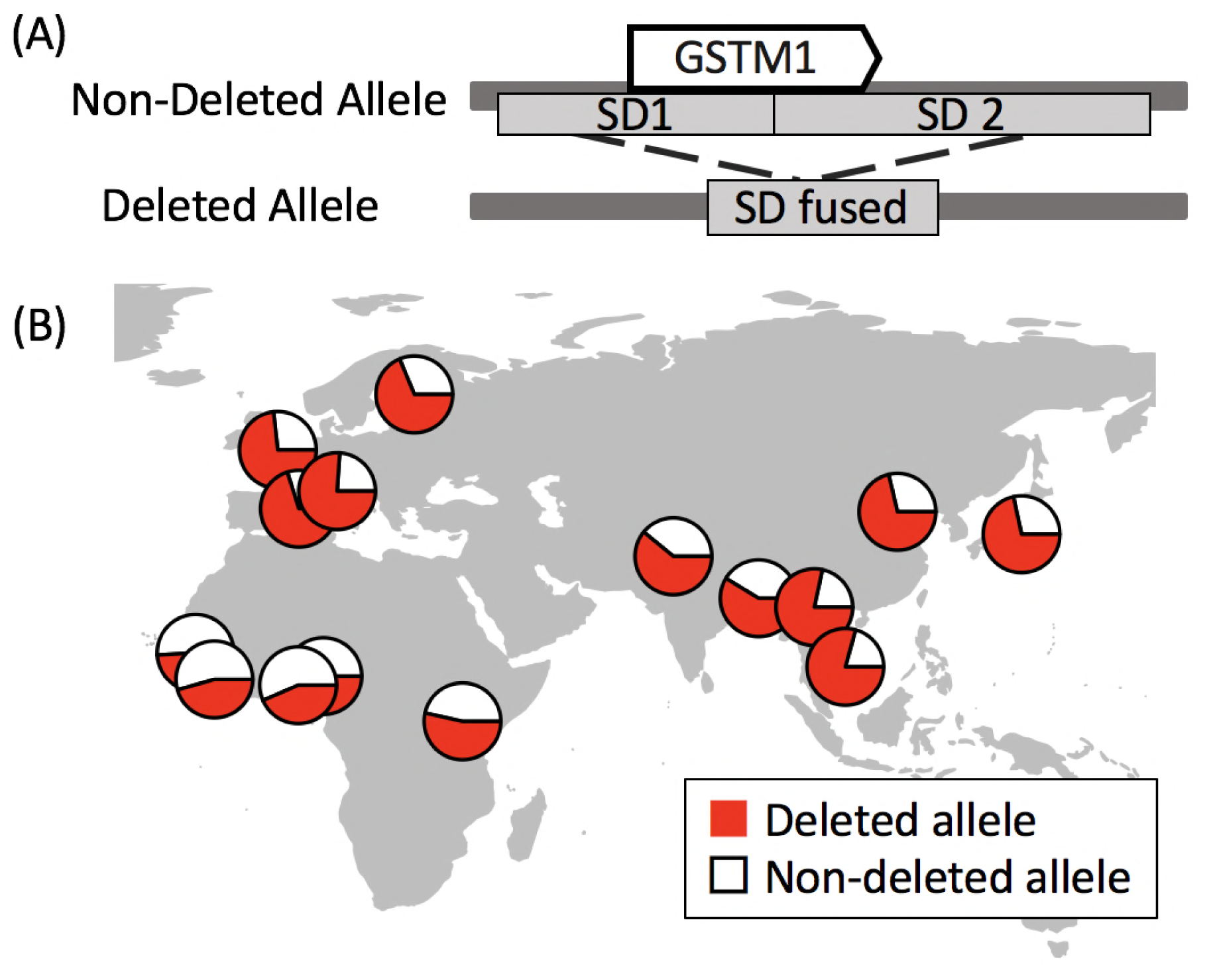
The formation of the *GSTM1* deleted allele and its geographical distribution. Two highly-similar segmental duplications fuse each other and form the deleted allele (upper). The geographical distribution of the *GSTM1* deletion allele (red) is shown in the lower figure. The frequency data is from the 1000 Genomes Project phase 3 database (Sudmant *et al*. 2015a).

Despite the biomedical importance of this locus, a clear understanding of the mechanism through which the *GSTM1* deletion and other variations in this locus actually affect phenotype remains elusive. We believe that understanding the underlying evolutionary forces that shape the variation in this locus will give us a framework to better target the functionally relevant haplotypes. Such an evolutionary analysis has been difficult since the *GSTM1* gene resides in a complex genomic region with two segmental duplications (SDs) flanking both sides of the gene (**Fig. 1A**). This particular genomic architecture predisposes *GSTM1* to non-allelic homologous recombination events, which can lead to gene conversion events and formation of new SVs. In fact, the polymorphic deletion observed in humans was almost certainly a result of such a mechanism (Xu *et al*. 1998). Moreover, we recently reported that recurrent deletions of the *GSTM1* gene have happened independently in the human and chimpanzee lineages and that multiple gene conversion events have defined genetic variation in this locus in primates (Saitou et al., in press). Even though the recurrence and gene conversion events are fascinating, it complicates the phylogenetic and broader evolutionary analysis immensely. This may explain the paucity of evolutionary analysis in the *GSTM1* locus, which is otherwise the target of more than 500 publications in the last 20 years, most of which are single-locus association studies implicating variation in this locus in susceptibility to multiple cancers (Parl 2005).

In addition to its biomedical relevance, the *GSTM1* gene deletion is one of the most common whole gene deletions observed among human populations (**Fig. 1B)** (Garte *et al*. 2001; Gaspar *et al*. 2002; Saadat 2007; Buchard *et al*. 2007; Fujihara *et al*. 2009; Piacentini *et al*. 2011). A whole gene deletion leads to the lack of the corresponding protein and thus it is thought to be deleterious in most cases. Therefore, it is rare for whole gene deletions to be at high frequencies (Conrad *et al*. 2010; Sudmant *et al*. 2015b). Nevertheless, the average global frequency of Eurasians with homozygous *GSTM1* deletion is higher than 50% (Sudmant *et al*. 2015a), which puts it among the most frequent whole gene deletions observed in Eurasians according to 1,000 Genomes Project database, the other being deletions affecting the *LCE3BC* (Late Cornified Envelope 3B and C) and the *UGT2B17* (uridine diphosphoglucuronosyltransferase), both of which have been shown to be evolving under non-neutral conditions (Xue *et al*. 2008; Pajic *et al*. 2016).

Collectively, the functional relevance of the *GSTM1* deletion, its mechanistic complexity, and its high frequency make it suitable as a model for studying the evolution of metabolizing gene families. Therefore, in this paper, we analyzed hundreds of genomes to resolve the haplotypic variation that defines the *GSTM1* deletion.

## Methods

### Study Populations

We primarily used data from 3 populations, YRI (Yoruba in Ibadan), CEU (Utah residents with Northern and Western European ancestry), and CHB (Han Chinese in Beijing) in the 1000 Genome Project (McVean *et al*. 2012; Sudmant *et al*. 2015b). For the phylogenetic analysis, we also analyzed haplotypes of the chimpanzee reference genome (The Chimpanzee Sequencing Consortium 2005) and the Neanderthal genome (Prüfer *et al*. 2014) and Denisovan genome (Reich *et al*. 2010).

### Confirmation of the genotyping and phasing in our dataset

To validate the accuracy of the genotyping of the *GSTM1* deletion in 1KGP dataset, we used digital droplet polymerase chain reaction experiments to genotype the *GSTM1* deletion in 17 humans from 1KGP (forward primer: TCGAGGGTGCCATTACATTC; reverse ACTTCTGTCCTGGGTCATTC; probe: /56-FAM/TAGGAGCAG /ZEN/GCAGGTGATGTGAAC /3IABkFQ/). We followed standard protocol provided by Bio-Rad EIF2C1 probe assay. The results were in **Table S1**.

To investigate the accuracy of the 1KGP genotyping of single nucleotide variants, we investigated 74 CHB individuals and 6 sites in the target regions (chr1:110211681-110223007 and chr1:110246810-110255596) reported both in 1KGP and HapMap (this is all the SNPs genotyped in HapMap in the region). Of all the 444 sites, 9 were NN in hapmap and 3 showed different results in the two databases. Overall, we found approximately %99.3 concordance between the two datasets.

To validate the accuracy of the phasing in 1KGP project data that we used in this study, we compared haplotypes on 50kb on each side of the deletion in our dataset with phased HapMap genomes. Specifically, we were able to find 322 heterozygous sites in 30 phased HapMap CEU genomes. 320 (∼99.4%) of these are concordant with the 1KGP dataset.

### Linkage disequilibrium and haplotype analyses

Vcftools (Danecek *et al*. 2011) was used to calculate R^2^ values between the *GSTM1* deletion and flanking single nucleotide variations in Han Chinese in Bejing, China (CHB), Utah residents with Northern and Western European ancestry (CEU) and Yoruba in Ibadan, Nigeria (YRI) populations in 1KGP phase 3 datasets (**Fig. 2A, Fig. S1**). The Ensembl genome browser (http://asia.ensembl.org/index.html) was used to calculate and visualize linkage disequilibrium (LD) between single nucleotide variations and between the deletion and single nucleotide variations. Based on the R^2^ values, we used two target regions for the statistical neutrality tests. Two target regions were selected for the following analyses; target1, upstream the deletion (chr1:110211681-110223007 in the hg19) and target2, downstream the deletion (chr1:110246810-110255596 in the hg19), which showed relatively high R^2^ with the *GSTM1* deletion in CHB **(Fig. 2A)**. We constructed a phylogenetic tree of the combination of these two regions by VCFtoTree (Xu *et al*. 2017a) and visualized it in Dendroscope (Huson *et al*. 2007). For the clustering and the visualization of haplotypes and its cluster analysis, we used the pipeline described in Xu et al., (Xu *et al*. 2017c). The clustering was conducted based only on the haplotypes themselves with neither *a priori* input of haplogroups nor the deletion status (**Fig. 3**).

**Figure 2.**
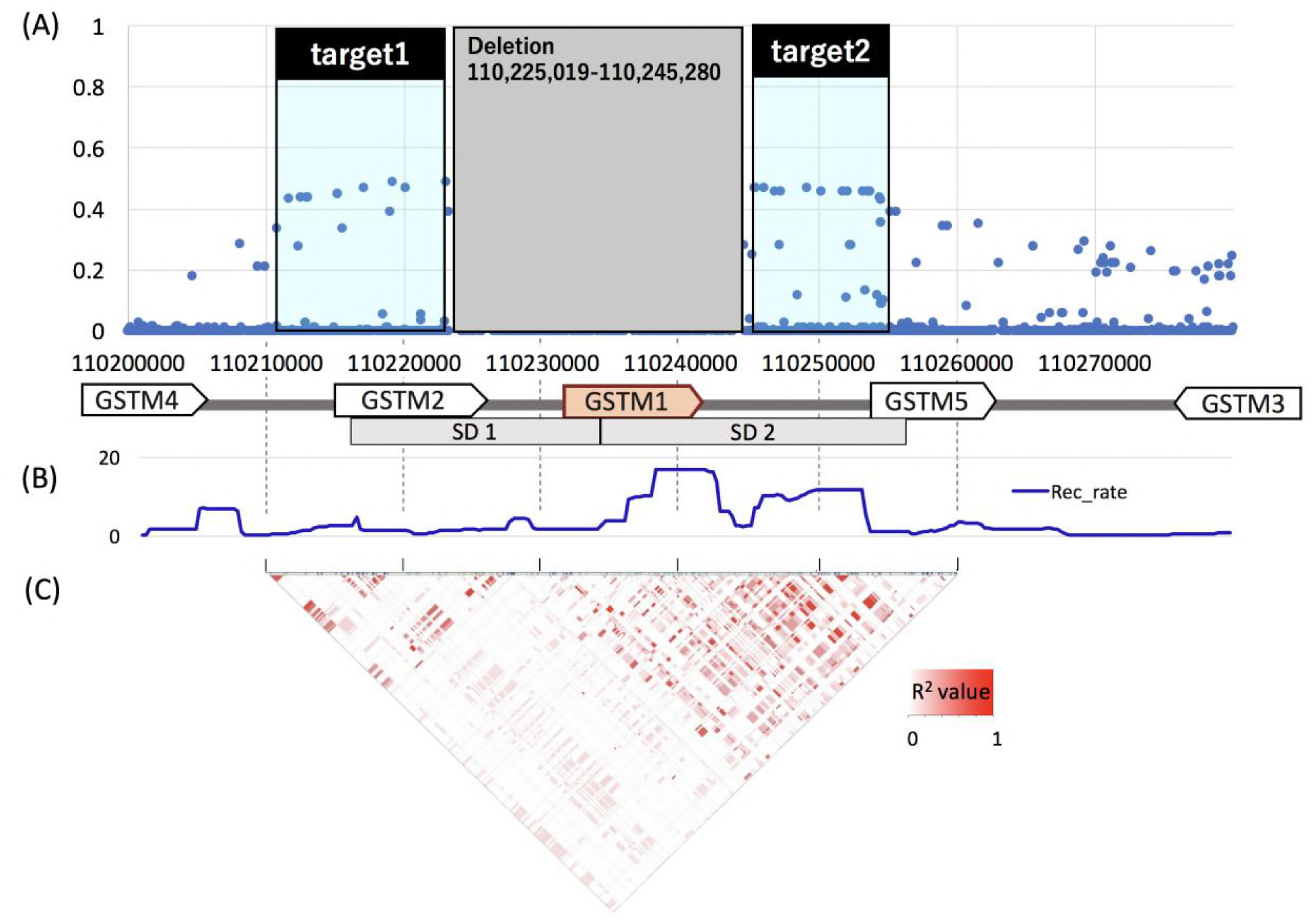
Linkage disequilibrium in the *GSTM1* region in CHB. (**A**) R^2^ value between the *GSTM1* deletion and flanking single nucleotide variants in CHB. Each dot represent each single nucleotide variants. X-axis indicates chromosomal location and Y-axis indicates the R^2^ value between the *GSTM1* deletion and flanking single nucleotide variants. Target regions were chr1:110211681-110223007 (target1, upstream the deletion) and chr1:110246810-110255596 (target2, downstream the deletion). (**B**) Recombination rate of the *GSTM1* flanking region.(Dib *et al*. 1996; Broman *et al*. 1998; Kong *et al*. 2002) (**C**) Linkage disequilibrium (LD, R^2^ value) between the single nucleotide variants around the *GSTM1* gene in CHB. Color coded in a continuous R^2^ values between each single nucleotide variants from 0 (white) to 1 (red).

**Figure 3.**
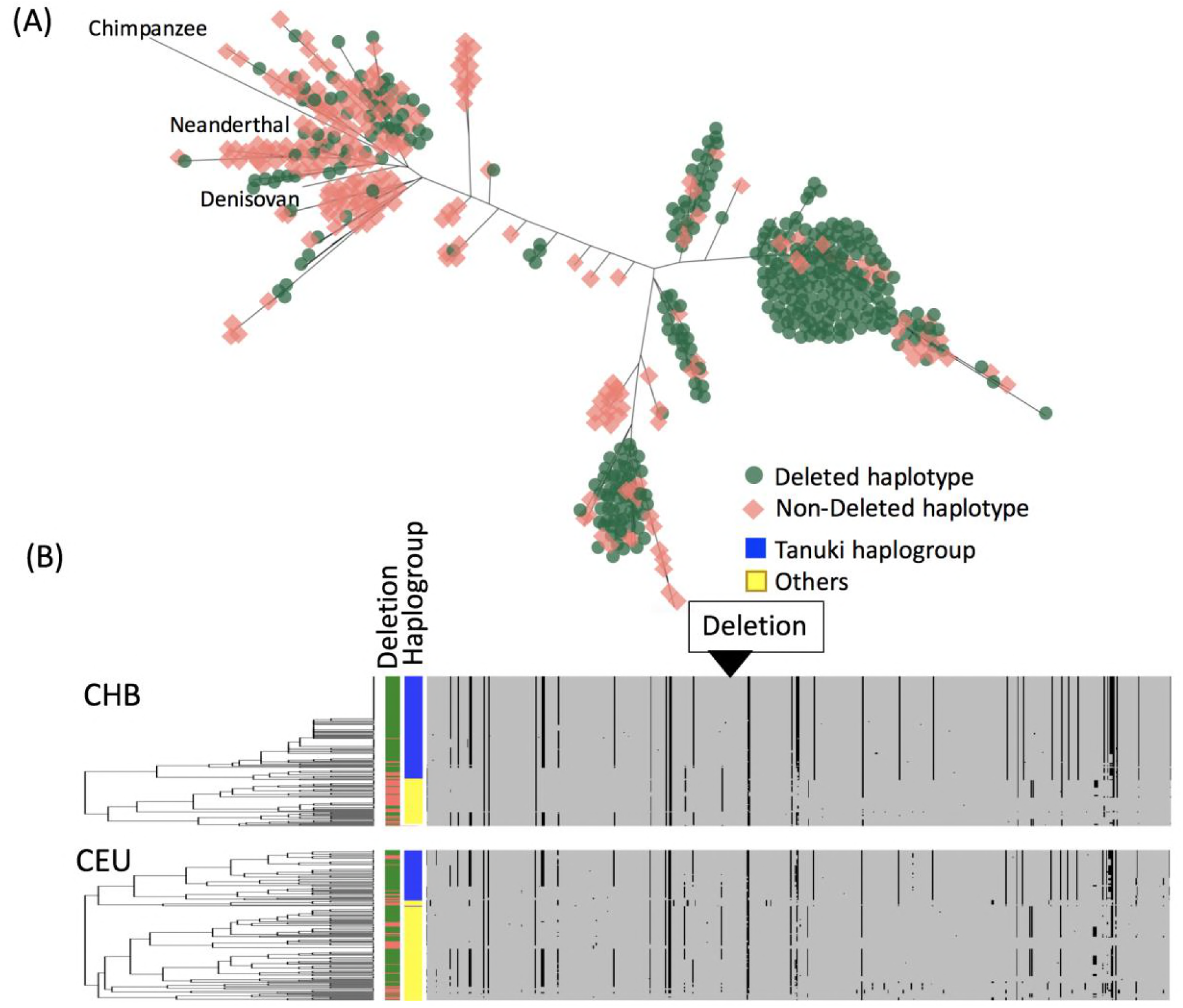
Haplotypes of the combined target1 and target2 regions. (A) The phylogenetic relationship of the downstream target 2 region of CEU, YRI, CHB, Neanderthal, Denisovan and Chimpanzee constructed by the maximum-likelihood method. The *GSTM1*-deleted haplotypes were marked by green and the *GSTM1*-non-deleted haplotypes were marked by red manually. (B) Clustered haplotypes of the upstream target 1 region and downstream target 2 regions of CHB and CEU. The clustering was conducted based only on the haplotypes themselves with neither *a priori* input of haplogroups nor the deletion status. The *GSTM1*-deleted haplotypes were marked by green and the *GSTM1*-non-deleted haplotypes were marked by red. The *Tanuki* haplogroup, which showed high population differentiation, was marked by blue and other haplotypes were marked by yellow.

**Figure 4.**
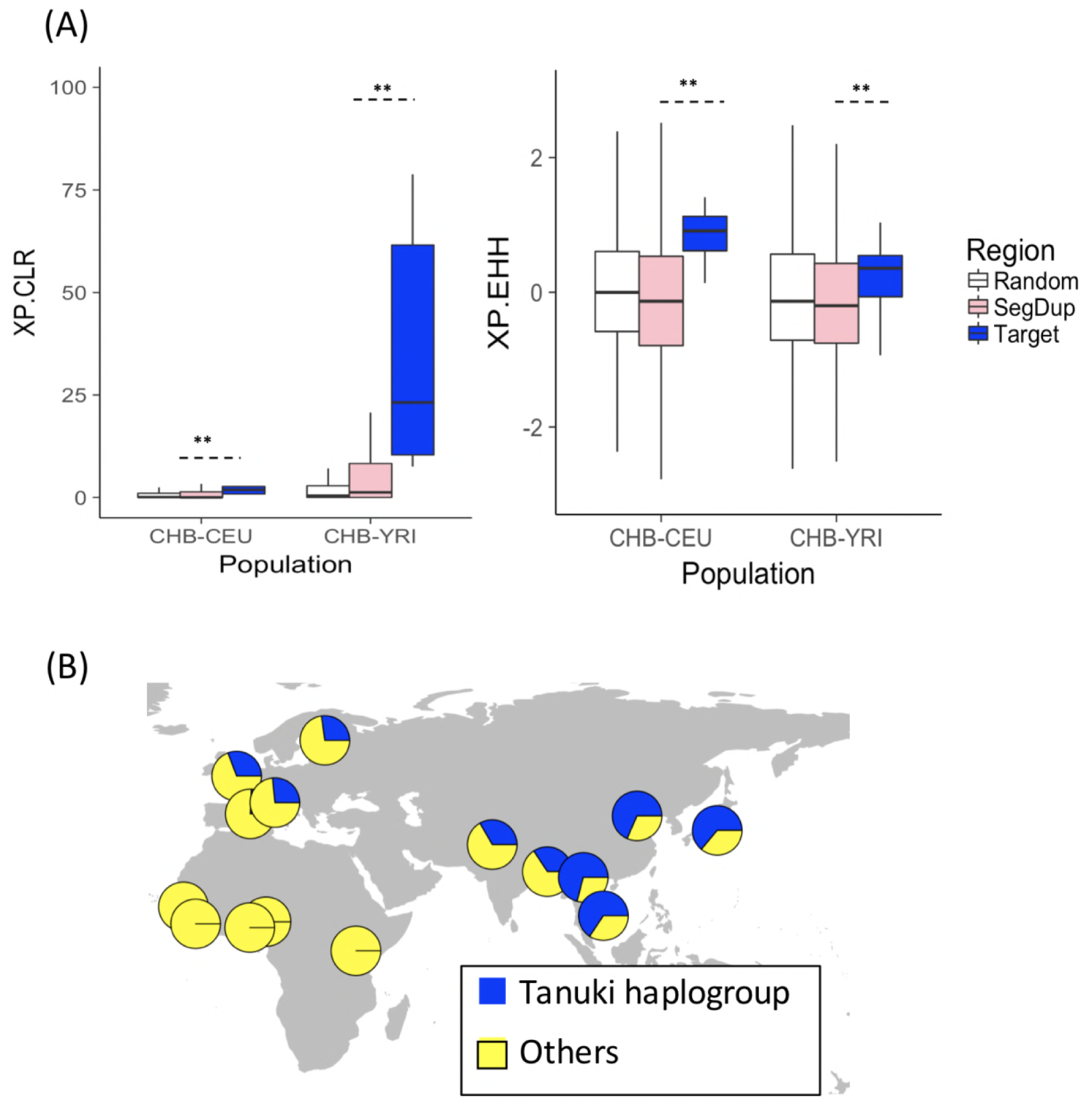
**(A)** Neutrality tests on the target region (chr1:110246810-110255596, downstream the deletion, also represented in Figure. 2.), the 1000 randomly selected 9kb regions, and all the segmental duplications on chromosome one, which contains the *GSTM1* in CEU, CHB and YRI populations in 1KGP. XP-CLR and XP-EHH were calculated for 2kb intervals. The stars show significant differences (p<0.01, Wilcoxon Rank Sum Test). **(B)** The geographical distribution of the *Tanuki* haplogroup calculated from the 1KGP phase3 data.

### Tests of natural selection on the Target region

We first constructed a null distribution by calculating the *F*_*ST*_ *(Weir and Cockerham 1984)*, Tajima’s D (Tajima 1989), π (Nei and Li 1979) and iHS (Voight *et al*. 2006) values for 1,000 randomly chosen regions on the same chromosome where the *GSTM1* is located (chromosome 1) and also match our target region by size (∼9kb). Then, we calculated these statistics observed in the downstream regions of the *GSTM1* deletion (chr1:110246810-110255596, downstream the deletion, also represented in **Fig. 2**) and compare the results with those from the random regions. We also replicated this with size-matched regions that overlap with all 618 segmental duplications on chromosome 1 to match the genomic features in the *GSTM1* region (**Fig. S2**).

We used genomic data of CEU, CHB and YRI populations in 1KGP. Tajima’s D value, iHS and π were calculated for 3kb intervals. XP.CLR were calculated for 2kb intervals. All these measurements were acquired through the 1000 Genomes Selection Browser (Pybus *et al*. 2014).

To visualize the geographic distribution of the *Tanuki* haplogroup, we used data from 15 populations in the 1000 Genome Project for haplotype distribution analysis: BEB, CDX, CHB, ESN, FIN, GBR, GWD, IBS, JPT, KHV, LWK, MSL, PJL, TSI, and YRI, which have not experienced recent population admixture or migration. We used the “rworldmap” package (South 2011) to visualize the global distribution of the haplotypes.

### Simulations for *F*_*ST*_ and π values

For the simulation analyses, we used ms (Hudson 2002) to conduct 1000 simulations matching the size (8,787bp) and the observed Watterson Estimator (θ=9.14) which was calculated by DnaSP (Librado and Rozas 2009) of the downstream target2 region **(Fig. 2)**. For each simulation, we generated 216 and 206 haplotypic sequences for YRI and CHB, respectively. We chose these populations and the number of sequences to match the empirical data from the 1000 Genome Project. To accurately model the effect of the demography in our simulations, we used the published parameters for East Asian and African populations (Schaffner *et al*. 2005) and we implemented the recombination rates of this region (ρ=5) reported by the HapMap Consortium (International HapMap Consortium *et al*. 2007). Once the sequences are generated, we focused on constructing expected distributions for population differentiation (*F*_*ST*_) and nucleotide diversity (π). As mentioned above, we avoid linkage based approaches, given the high levels of gene conversion in the locus.

The command line for ms simulator was ./ms 422 1000 -t 91.4 -I 2 216 206 -m 1 2 3.2 -n 2 0.077 -en 0.0005 1 0.25 -en 0.001 2 0.077 -ej 0.00875 2 1 -en 0.0425 1 0.125 -r 5 8787.

We calculated *F*_*ST*_ values between YRI and CHB populations for each single nucleotide variant that was generated in the simulations. We then plotted the values with the matching allele frequencies (0.69-0.70, the range of standard deviation of the observed allele frequency of the variations in *Tanuki*) for each of these variants. Next, we considered the distribution of the simulated π values. To simulate this, we first generated 1,000 simulated datasets as described above. We then focused on the segregating sites with alternative allele frequencies at 0.69-0.70, the range of standard deviation of the observed allele frequency of the variations in *Tanuki* haplogroup. Based on this, we calculated π for each of these haplogroups which carry the alternative allele for each single nucleotide variation in the simulated CHB population (**Fig. S3**).

To further explore this issue, we have considered the π values for *Tanuki* and *nonTanuki* haplotypes across a larger genomic region surrounding the *GSTM1*, including sequences that are not overlapping segmental duplications in multiple East Asian populations (CHB, CHS, JPT and KHV) (**Fig. S4**). These results also corroborates our initial finding that *Tanuki* haplotypes show unusually low π, as compared to simulated expectations (**Fig. 5B**). Acknowledging that other possibilities that we did not consider may exist (e.g., such as effect of negative selection of nearby genes), our current results are most parsimonious with the adaptive evolution of *Tanuki* haplogroup in East Asian populations.

**Figure 5.**
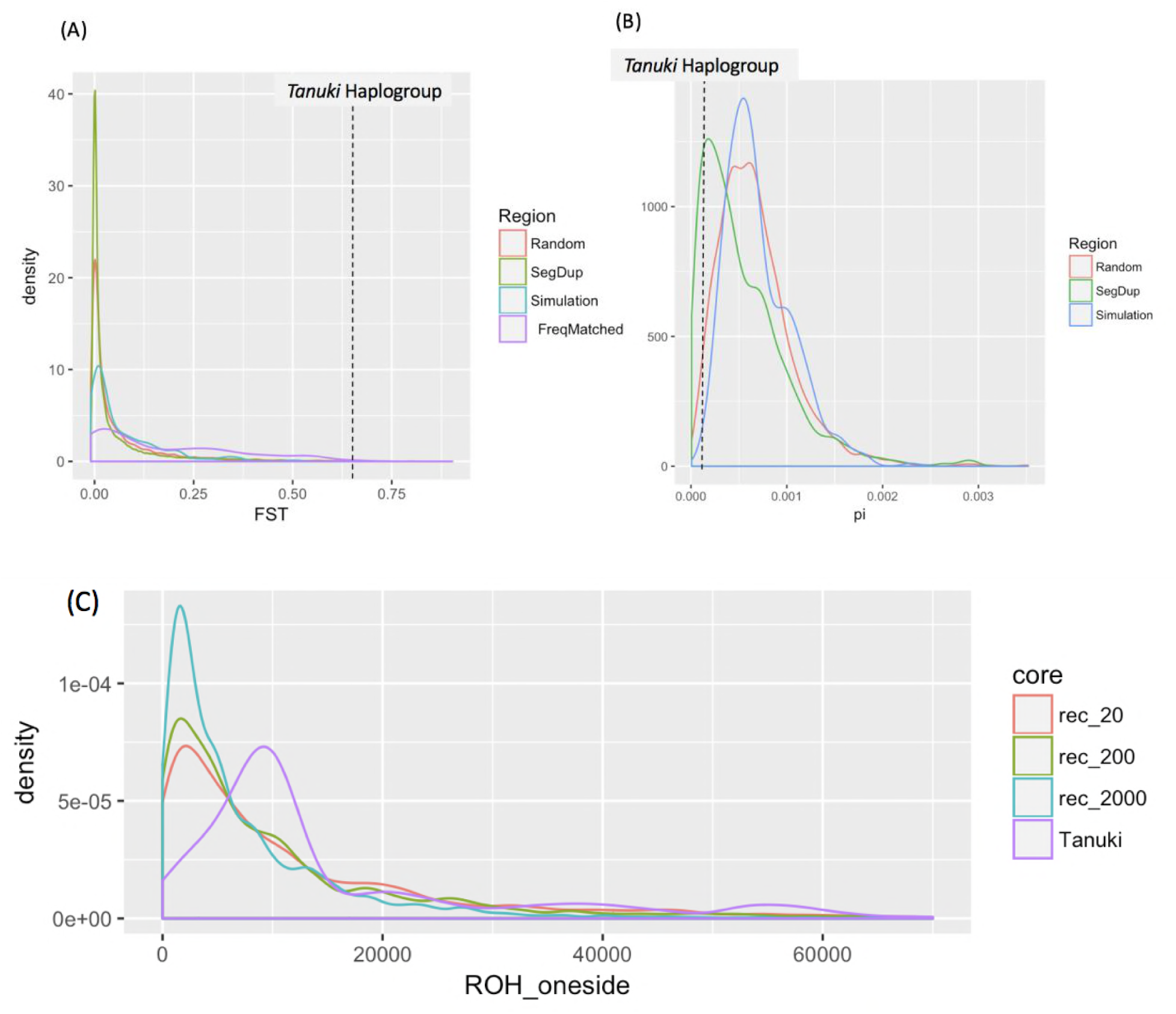
(**A**) The average values of *F*_*ST*_ for *Tanuki* SNPs (dotted line), the 1000 randomly selected 9kb regions (red), 9kb regions on chromosome 1 that overlap with segmental duplications (green), simulated data under neutrality (blue), and frequency-matched (0.69-0.70) single nucleotide variants on chromosome 1 (purple). (**B**) The π value for *Tanuki* haplotypes (dotted line), the 1000 randomly selected 9kb regions (red), 9kb regions on chromosome 1 that overlap with segmental duplications (green), simulated data under neutrality (blue). (**C**) The Run of Homozygosity of the *Tanuki* SNP (purple) and simulated frequency-matched SNPs with 3 different recombination rates, 20 (red), 200 (green), 2000 (blue).

### The Run of Homozygosity analysis

We used the demographic model of (Schaffner *et al*. 2005) with theta=0.00375 per site and Rec rate=0.001 per site and 4Nem=9 from the empirical observation. We ran more than 7000 simulations per each condition and obtained 70-120 core sites with the frequency of 0.69+-0.031 (Standard deviation calculated from Binomial Distribution) in East Asia and 0 in Africa under neutrality. We calculated the both side of ROH for the 70-120 core sites from the simulation. We also calculated one side (downstream) of ROH of the 238 *Tanuki* homozygous individuals in East Asian populations for the *Tanuki* SNP as there was a recombination hotspot in the upstream of *Tanuki* region (**Fig. 2B**). We used three recombination rates (20, 200, 2000 recombinations per the 200kb region) to see the impact of recombination rates on ROH. r=200 was the most realistic parameter from the observation.

The command line for ms simulator was ./ms 422 1000 -t 750 -r 200 200000 -I 2 206 216 9 –en 0.0005 2 0.25 -en 0.001 1 0.077 -en 0.0425 2 0.125

To test for potential sampling bias (*e*.*g*., consanguinity) in our dataset, we conducted bootstrap resampling of individual phased chromosomes. There are 238 individuals in East Asian populations (CHB, JPT, CHS, KHV, CDX) that are homozygous for *Tanuki* haplotype. To construct a distribution to investigate whether there is a sampling bias, we randomly resampled from 694 phased Tanuki haplotypes to generate 500 datasets each with 238 homozygous constructs. This allowed us to test whether the observed value of ROH deviates from randomly resampled distribution. We observed no such deviation, suggesting no sampling bias in our analysis (**Fig S5**).

### Age estimation of *Tanuki* haplogroup and selection on *Tanuki* haplogroup

Assuming a divergence time of 6 million between chimpanzee and humans, we estimated the coalescent of *Tanuki* haplogroup (0.001048 distance from the root haplotype) to be approximately 360 thousand years before present. At the past time when r*L*t reaches 1, the ROH shows rapid breaks because of recombination (Racimo *et al*. 2015). So, we can assume that the t at that time is the start point of the selection. We used r= 1×10^−8^/generation, u=3×10^−8^/site/generation, and observed averaged ROH_aver_ of the *Tanuki* Haplogroup, 16kb, and estimated T=1500 generations=46K years ago when the selection on *Tanuki* Haplogroup began.

## RESULTS

### Imperfect linkage disequilibrium between the *GSTM1* deletion and flanking single nucleotide variants indicate multiple gene conversion events in the human lineage

To understand the evolutionary mechanisms which maintain the *GSTM1* deletion common in humans, we first attempted to resolve the haplotype structure of the locus. We calculated the R^2^ value between the *GSTM1* deletion and flanking single nucleotide variants in Han Chinese in Bejing, China (CHB), Utah residents with Northern and Western European ancestry (CEU) and Yoruba in Ibadan, Nigeria (YRI) populations in 1KGP phase 3 datasets (**Fig. 2A, Fig. S1**). Remarkably, we found almost no linkage disequilibrium between the *GSTM1* deletion flanking single nucleotide variants in CEU and YRI (**Fig. S1**), and observable, but imperfect linkage disequilibrium in CHB only extending to >2.5kb on each side of the deletion (R^2^=∼0.45, **Fig. 2A**). One way to explain this low level of linkage disequilibrium is by invoking a recombination hotspot in the region. However, we found that the reported recombination rate for the region is high, but not exceptionally high to be categorized as a hotspot (3-17 cM/Mb,(Mc Vean *et al*. 2004)) (**Fig. 2B**).

We reasoned that the imperfect linkage disequilibrium between the deletion and the neighboring variants can be due to two mechanisms: (i) recurrent formation of deletions or (ii) exchange of genetic material across chromosomes because of gene conversion events. One way to test this was described by Boettger et al. (Boettger *et al*. 2016). Specifically, they focused on the linkage disequilibrium between the variants on each side of the deletion. Note here that it is already known that there is weak linkage disequilibrium between the deletion and the flanking variants. The question is whether the flanking regions have linkage disequilibrium with each other, especially when haplotypes with the deletions are considered. If they do, this would indicate that the deletion is independently formed more than once (*i*.*e*., recurrent evolution), breaking the linkage disequilibrium between the deletion and the flanking regions, without disturbing the linkage disequilibrium between the flanking regions themselves. In contrast, a gene conversion event would break the linkage disequilibrium between the flanking regions on each side of the deletion. We tested these two scenarios and found that there is weak linkage disequilibrium between variants on each side of the deletion in CHB (**Fig. 2C**), supporting the scenario of interlocus gene conversion in this locus. In addition, we have already found multiple gene conversion events among different primate lineages affecting this locus (Saitou et al. In Press). However, the occurrence of gene conversion is not mutually exclusive with the recurrence of the deletion. As such, it is plausible that recurrent deletions may have occurred in this locus in addition to the gene conversion events. Regardless, it is clear from our analysis that gene conversion is a major process shaping the linkage disequilibrium patterns in this locus.

To further understand the haplotypic variation in this locus, we constructed a phylogenetic tree using sequence variation data in the “target” region flanking the deletion for 620 phased haplotypes from YRI, CEU, CHB populations, as well as chimpanzee, Denisovan, Neanderthal genomes (**Fig. 3A**). It is important to note that we have confirmed and validated both the accuracy of deletion genotypes and phasing in 1,000 Genomes dataset using digital PCR-based genotyping and array-based HapMap database, respectively (see Methods). As expected from the low levels of linkage disequilibrium between single nucleotide variants and the deletion, we observed no clear separation between haplotypes with and without the deletion. Instead, we found multiple branches that are predominantly populated with deleted haplotypes and others with non-deleted haplotypes without notable population structuring. It is important to note here, however, that Neanderthal and Denisovan haplotypes, as well as the branching point for the chimpanzee reference haplotype, all cluster with the predominantly non-deleted haplotypes. This suggests that the deletion is a derived variant that likely evolved after the human-Neanderthal divergence (i.e., in the last 1 Million years).

Despite these general insights, the observable lack of linkage disequilibrium between the *GSTM1* deletion and the neighboring variants reduces the power of disease-association studies (*e*.*g*. GWAS) that rely on imputation of the *GSTM1* deletion genotype using nearby single nucleotide variants. Even considering multiple single nucleotide variants within a single population, we were not able to impute the deletion accurately in any of the study populations (YRI, CHB, and CEU) (**Fig. 3B**). This is concordant with the previous finding reported the difficulty to predict the copy number of the *GSTM1* gene by the flanking haplotypes in CEU (Khrunin *et al*. 2016). As such, direct genotyping, rather than imputation, may be more robust approach to study *GSTM1* deletion. In sum, our analyses suggest that the variation in the *GSTM1* locus in humans has undergone multiple gene conversion and recombination events, complicating the haplotype architecture associated with the deletion.

### A potential signature of a soft-sweep in East Asian populations in the *GSTM1* locus

To reveal any potential signatures of adaptive evolution affecting variation in this locus, including the *GSTM1* deletion, we conducted statistical neutrality tests using single nucleotide variation data from the sequences flanking the *GSTM1* deletion. Specifically, we first constructed a null empirical distribution by calculating the *F*_*ST*_ *(Weir and Cockerham 1984)*, Tajima’s D (Tajima 1989), π (Nei and Li 1979), iHS (Voight *et al*. 2006), XP-EHH and XP-CLR values for 1,000 randomly chosen regions on the same chromosome where the *GSTM1* is located (chromosome 1) and also match our target region by size (∼9kb). Then, we calculated these statistics observed in the downstream regions of the *GSTM1* deletion and compared the results with those from the random regions (**Fig. 4A, Fig. S2**). We also replicated this with randomly selected, size-matched regions that overlap with all of the 618 segmental duplications on chromosome 1 to match the genomic features in the *GSTM1* region (**Fig. S2**). We found that both XP-CLR and XP-EHH tests showed significant differences as compared to neutrality when *CHB* population is involved (*p* values<0.01, Wilcoxon rank sum test with continuity correction). These tests measures the change in allele frequency in one population has occurred more quickly than expected by drift alone and the difference in extended homozygosity between two populations (**Figure 4A, Table 1**). XP-EHH (Sabeti *et al*. 2007) is cross-population extended haplotype homozygosity and XP-CLR (Chen *et al*. 2010) is multi-locus allele frequency differentiation between two populations. Collectively, the unusually high XP-CLR and XP-EHH values in CHB population suggest a incomplete soft sweep has shaped the distribution of *Tanuki* haplogroup in this population.

**Table 1.**
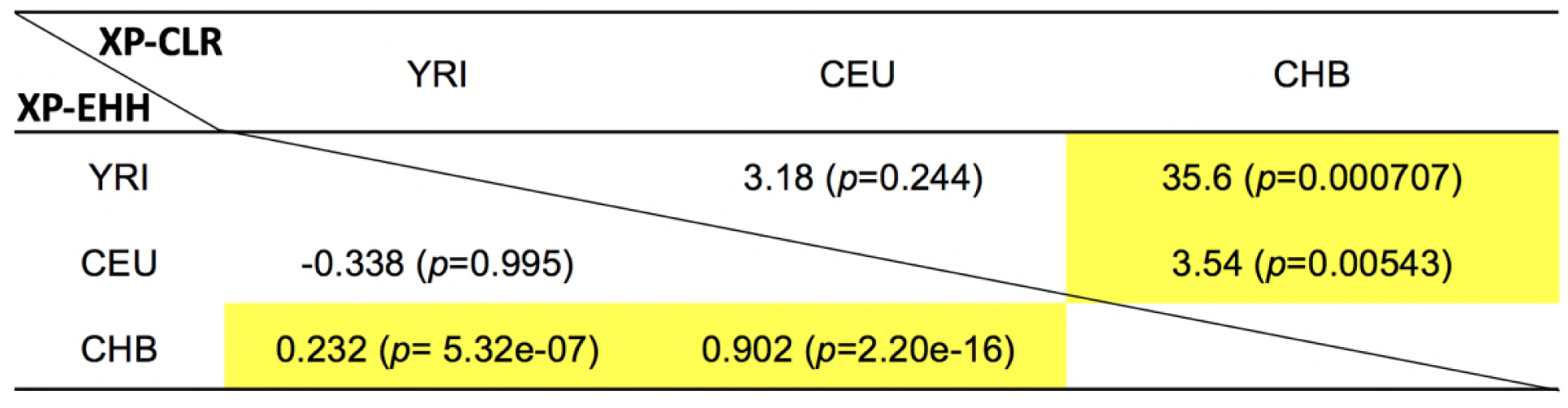
XP-EHH and XP-CLR between CHB-YRI, CEU-YRI and CHB-CEU. Significant values were highlighted yellow.

To investigate the haplotypic background of this putative soft sweep, we focused on the single nucleotide variants that showed the highest population differentiation (**Fig. 4A**). We first showed that they are in high linkage disequilibrium with each other (R^2^>0.84), indicating that they represent a single haplotype group in East Asian populations (**Fig. S6, Table S2**, called *Tanuki* haplogroup from hereon) in the target2 region (**Fig. 2A**). We first investigated the global distribution of the haplogroup and found that it represents the major allele (>70% allele frequency) in East Asian populations, but it is virtually absent in sub-Saharan African populations **(Fig. 4B, Fig. S7)**.

To determine the age of this haplogroup, we constructed a phylogenetic tree (**Fig. S8)** of all *Tanuki* haplotypes in CEU and CHB populations (no *Tanuki* haplotype was found among YRI genomes), along with a non-*Tanuki* haplotype and the chimpanzee reference genome haplotype as outgroups. Then, assuming a divergence time of 6 million between chimpanzee and humans, we estimated the coalescent of *Tanuki* haplogroup to be approximately 360 thousand years before present. Even considering the relatively large standard deviation of the estimate, this date is far earlier than the out-of-Africa migrations. As such, we conclude that this haplogroup has originated in Africa, but has increased in frequency in Eurasia. It is plausible that some *Tanuki* haplotypes still remain in African populations, but at a low allele frequency.

To further understand the haplotypic background of the significantly high XP-CLR and XP-EHH values in Asian populations as compared to European and African populations, we constructed a network of *Tanuki* haplotypes (**Fig S9**). We found that a vast majority of *Tanuki* haplotypes in CHB population has identical sequences, suggesting a rapid and recent increase in the allele frequency in the East Asian populations. This observation explains the unusually high XP-CLR and XP-EHH values observed for CHB population and further support the notion that a incomplete soft sweep has shaped the distribution of *Tanuki* haplogroup in this population.

### Allele frequency of the *Tanuki* haplogroup is unusual in East Asia, but not in Europe

The high XP-CLR and XP-EHH values along with very high frequency of *Tanuki* haplogroup in CHB potentially point to the effect of an local soft-sweep. The alternative, null hypothesis would be that the observed difference in allele frequency is due to the effect of drift alone. To further distinguish between these two scenarios, we conducted multiple empirical and simulation based analyses. First, we investigated whether the allele frequency of *Tanuki* haplogroup in East Asian and European populations can be explained by drift alone. To do this, we first assumed that the putative target for selection is an allele that reached to approximately 70% allele frequency in East Asia, but remain at 25% allele frequency in Europe (**Fig. 4B**). Then, we used the simple formula detailed (15) in (Kimura and Ohta 1973) and estimated that it would take ∼580 and ∼160 thousand years under neutrality for that single haplotype to increase in allele frequency from 1/2N_e_ in ancestral African population to 70% and 25% observed in Asia and Europe, respectively. This results suggest that the allele frequency of *Tanuki* haplotypes in Europe can be explained by neutrality alone, but not in East Asia.

To further interrogate whether the high allele frequency of *Tanuki* haplotypes can be explained by genetic drift alone, we conducted simulation-based analyses of the genetic variation we observed (for detailed conditions for simulations, see Methods). Specifically, we used ms (Hudson 2002) to generate 1,000 simulated datasets comprising sequences matching the size (8,787bp) of the target2 region of *GSTM1* **(Fig. 2)**. For these simulations, we used demographic parameters previously laid out for CHB and YRI populations (Schaffner *et al*. 2005) to accurately model the effect of drift on nucleotide diversity and allele frequency.

Once the sequences were generated, we calculated *F*_*ST*_ values for each single nucleotide variant that was generated in the simulations between two simulated populations (YRI and CHB). To verify the accuracy of our simulations, we first compared our simulated sequences to empirical distribution of *F*_*ST*_ between YRI and CHB populations for randomly selected single nucleotide variants across the genome (**Fig. 5A**). We then plotted the *F*_*ST*_ values with the matching allele frequencies (0.69-0.70, that represent the deviation of the observed allele frequency of the variations in *Tanuki* in Asian populations) for each of these variants observed in the simulation. We found that none of 603 frequency-matched simulated *F*_*ST*_ is higher than the observed *F*_*ST*_ for the *Tanuki* haplogroup **(Fig. 5A)**. Overall, these results suggest that the unusually high allele frequency of *Tanuki* haplotypes in CHB population is unlikely to be explained by neutral forces alone.

### Haplotype homozygosity provides further evidence for a soft-sweep

In parallel to the allele frequency based analyses described above, we conducted tests based on haplotype similarity. We first followed the reasoning outlined by Kim and Satta (Kim and Satta 2008). Briefly, if a recent soft-sweep indeed increased the allele frequency of a particular haplotype group, we expect that the nucleotide diversity would decrease among the haplotypes that were sweeped, but not the others. Indeed, we found that the nucleotide diversity of haplotypes that belongs to the *Tanuki* haplogroup, which make up >70% of haplotypes in East Asians, is approximately six times lower (π=0.00012) in East Asians as compared to the nucleotide diversity observed in haplotypes that do not belong to the *Tanuki* haplogroup in the same population (π=0.00076).

To quantify this observation, we used simulated sequences (as described above) to construct an expected, neutral distribution of nucleotide diversity (π) values. Specifically, we wanted to test whether π calculated for the haplotypes that belong to the *Tanuki* haplogroup is lower than expected distribution under neutrality. To simulate this, we first generated 1,000 simulated datasets as described above. From the simulated dataset, we chose haplogroups within the simulated with alternative allele frequencies at 0.69-0.70 to match that of *Tanuki* haplogroup in frequency (please see **Fig. S3)**. Then, we plotted the π values for these simulated haplogroups. The results showed that π observed in the *Tanuki* haplogroup is lowest among 389 frequency-matched simulated values (**Fig. 5B**).

To test whether the low nucleotide diversity observed in *Tanuki* haplogroup is also translated into extended homozygosity, a hallmark of a selective sweep, we simulated expected Runs of Homozygosity (ROH) assuming demographic parameters outlined in (Schaffner *et al*. 2005) and different recombination rates (see methods for details). We found that the ROH values observed for the *Tanuki* single nucleotide variants in the 1KGP East Asian populations is significantly higher than what is expected under neutrality based on frequency matched simulation results with r=200, the most realistic parameter from the observation (**Fig. 5C**, p<2.5×10^−12^, Wilcoxon-Rank-Sum Test). The averaged ROH of the *Tanuki* haplogroup was 16Kb.

This calculation also allowed us to calculate a potential date for the sweep using the approach outlined elsewhere (Racimo *et al*. 2015) (see methods). Specifically, we estimated that a sweep starting approximately 46 thousand years ago affecting the *Tanuki* haplogroup, may explain the observed pattern in present-day CHB population. Combined, the observed values of population differentiation and haplotype diversity are inconsistent with the neutral evolution of this locus and are parsimonious with our hypothesis that the *Tanuki* haplogroup has increased its frequency under adaptive evolution in the East Asian populations.

### Potential functional impact of *Tanuki* haplogroup

By carefully resolving the different haplotype groups in the *GSTM1* locus, we were able to detect a putatively adaptive haplogroup (*Tanuki* haplogroup). The exact underlying evolutionary reason and the functional impact of the increase in frequency of the *Tanuki* haplogroup remain excellent venues for future studies. One obvious question is whether the *GSTM1* deletion is the actual target of positive selection. Indeed, we discovered that *Tanuki* haplogroup is significantly, albeit imperfectly, linked with the *GSTM1* deletion in the CHB population (R^2^=∼0.47) (**Table S1**). It is important to note that despite the low linkage disequilibrium, ∼91% of the *Tanuki* haplotypes harbor the *GSTM1* deletion. We reasoned that if the *GSTM1* deletion was the target for the selective sweep that we observed, then other haplotypes carrying the *GSTM1* deletion also show evidence for selection. However, we did not observe any unusual increase in allele frequency of other haplotypes that carry the deletion. As such, it is highly plausible, as drastic an event as a whole gene deletion is, *GSTM1* deletion may not be the main target of the selective sweep that we observed.

To further follow up this thread, we asked whether the *GSTM1* gene has no fitness consequences and hence accumulating loss of function mutations, such as the deletion. Under this scenario, the *GSTM1* deletion just swept along with the remainder of the *Tanuki* haplogroup with scant adaptive consequences. Contradicting this possibility, it was reported that the *GSTM1* gene sequence is conserved among great apes, as well as between humans and archaic humans. Specifically, we found only 4 nonsynomous variants between human and chimpanzee *GSTM1*, and found none among human, Neanderthal and Denisovan *GSTM1* genes (**Table S2**). In addition, only two commonly observed nonsynonymous mutations were reported for *GSTM1* gene in human population, both of which have little effect on the *GSTM1* function (Moyer *et al*. 2007; Tatewaki *et al*. 2009). ExAC database allowed us to be more specific (Lek *et al*. 2016). The expected number of loss-of-function variants at the *GSTM1* locus is 8.4 but observed number was 2 in the Exome analysis in 60,706 humans. In addition, the frequency of these loss-of-function variants are extremely low (<0.00003133). It is important to note that the deletion of the entire gene in approximately half of human genomes is not currently reported in ExAC database. However, our point here is that if *GSTM1* has no fitness effect in humans, we expect to find relatively high frequencies of loss of function variants in addition to the deletion. Instead, virtually all the nondeleted *GSTM1* haplotypes carry an intact, open-reading frame, further supporting the notion that both deleted and nondeleted haplotypes are maintained in the population.

These results suggest that it is likely that the *GSTM1* deletion has fitness effects, but it is independent of the selective sweep we observed for the *Tanuki* haplogroup. To explain these observations, we hypothesize that *Tanuki* haplogroup affect the function of multiple genes in the *GSTM* gene family and not just *GSTM1*. Consequently, the putative adaptive phenotype is a result of combination of these effects. Such a perspective has precedent where other studies described likely adaptive haplotypes that have effects on multiple metabolism gene families, such as *IRX* (Claussnitzer *et al*. 2015) and *FAD* (Fumagalli *et al*. 2015). To test whether *Tanuki* haplogroup indeed affect expression levels of *GSTM* genes other than *GSTM1*, we used gene expression data from Gtex portal (Lonsdale *et al*. 2013; Wheeler *et al*. 2016). Our results showed that this haplogroup is associated with significant decreases in expression of the *GSTM5* gene in most tissues, especially in the brain, but increases the expression of *GSTM3* in skeletal muscle (**Fig. S10**). As such, one potential argument would be that the selected effect is not the functioning of a single gene, but the overall regulatory impact of the *Tanuki* haplogroup on the *GSTM* locus.

## DISCUSSION

### Working with *complex* genomic structural variation loci

Our results provide a case study for the evolutionary impact of common haplotypic variation in a complex locus, involving both functional single nucleotide and copy number variants. One particular challenge in this locus that we verified is the general lack of linkage disequilibrium between common, neighboring variants. Specifically, we found a lack of strong linkage disequilibrium between the *GSTM1* deletion and neighboring variants. This finding corroborates our previous work at a deeper evolutionary depth, where we found evidence for multiple gene conversion events differentiating human and chimpanzee *GSTM* locus (Saitou et al., under review). As such, most genome wide association studies would not have effectively interrogated the potential biomedical impact of the *GSTM1* deletion because they have only investigated the single nucleotide variants, which do not adequately tag the deletion variant. In fact, when we consider the best case scenario for such studies and focus on East Asia, where the linkage disequilibrium is relatively high (R^2^=0.4), statistical power for a single nucleotide variant based genome wide association study would still be low. For example, based on Hong and Park (2012)’s calculations, the statistical power to detect an association would be only 26.5% when assuming a relatively standard experimental setup (*e*.*g*., the odds ratio of the phenotype 1.3, 1000 cases and 1000 controls, *etc*.).

This is not an exceptional case since the majority of deletions reside in similar haplotypic architectures (Sudmant *et al*. 2015b), complicating both evolutionary analysis, as well as disease association studies that depend on imputation to interrogate structural variants. Almost one-third of the deletions reported in 1000 genomes have R^2^<0.6 with neighboring variants (Sudmant *et al*. 2015b). As such genome-wide association studies will lose significant amount of power when assessing the effect of such SVs on tested trait if they are using imputation-based genotyping. Consequently, it is important to conduct direct genotyping of the deletion for such analyses or carefully resolve the haplotype structure within the locus to accurately assess the biomedical impact of these variants. Such locus-specific analyses indeed led to important connections between several structural variants and human diseases (Rothman *et al*. 2010; Usher *et al*. 2015; Boettger *et al*. 2016).

### Evolution of the *GSTM1* locus

Here, we report a soft sweep that increase the allele frequency increase of a particular haplogroup (*Tanuki*) in East Asian populations, putatively affecting the function of multiple *GSTM* genes (as summarized in **Fig 6**). Recent studies have shown the importance of soft sweeps on standing variation (Schrider and Kern 2017), rather than hard sweeps on novel variants (Hernandez *et al*. 2011), to be the dominating type of positive selection acting on human genome. In other species, such as Birch, soft sweeps were discussed as a major force specifically maintaining copy number variants of gene families (Salojärvi *et al*. 2017). In humans, ancient genic deletions and local structural variants have been discussed as possibly being maintained by such soft sweeps (Iskow *et al*. 2012; Lin *et al*. 2015).

**Figure 6.**
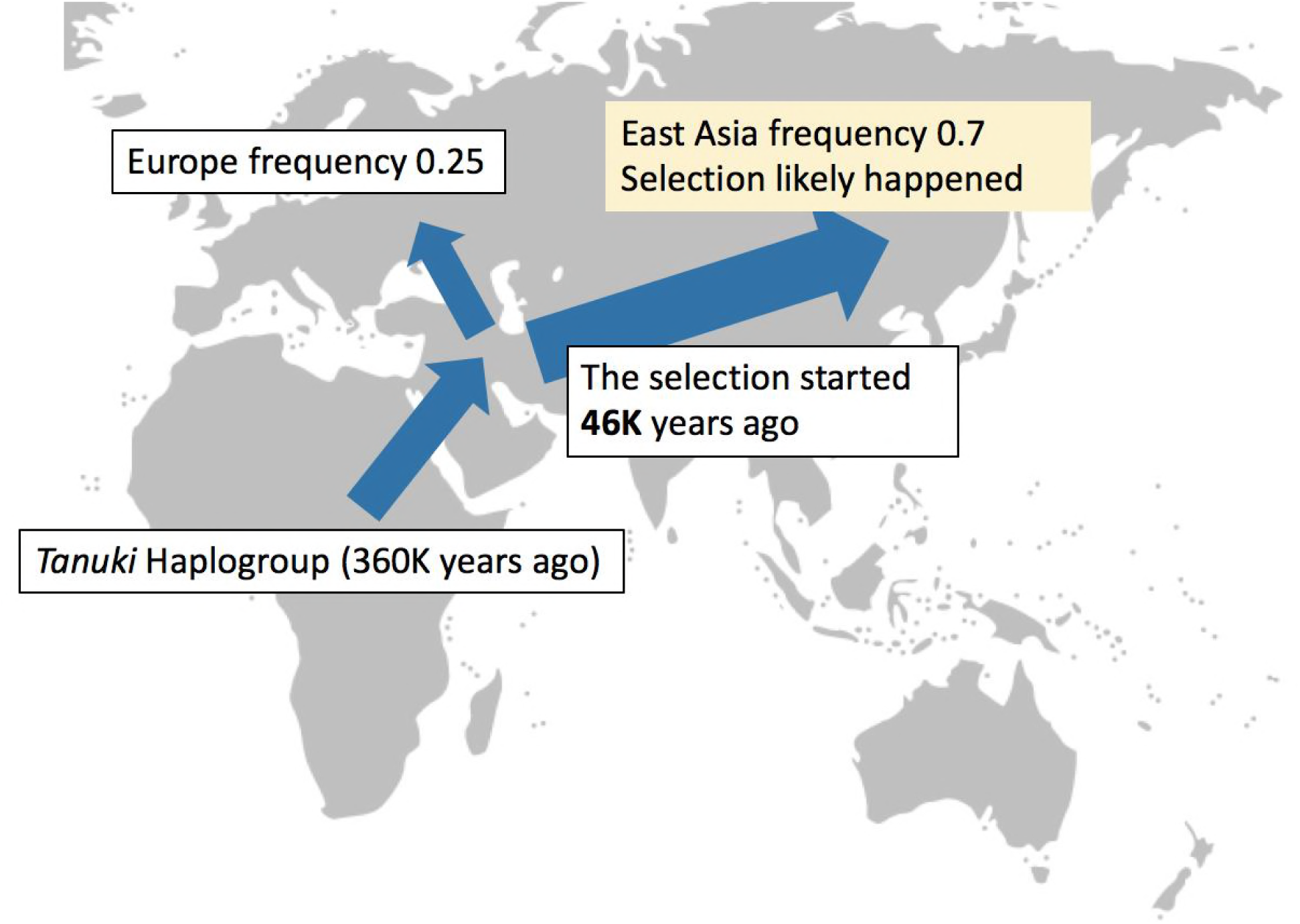
The evolutionary history of the *Tanuki* haplogroup. Based on our analyses, we argue that the *Tanuki* haplogroup originated before Out of Africa migrations (∼360K years before present) and spread out to Eurasia. We further argue that in Europe, the frequency of Tanuki haplogroup increased approximately to 0.25 under neutrality. In Asia, the selection on the *Tanuki* haplogroup started 46K years ago and pushed frequency of the *Tanuki* haplogroup to 0.7 under selection.

These insights fit well with our observations for the *Tanuki* haplogroup as well as with the emerging, broader picture where genes involved in metabolism have been shown to evolve under complex adaptive forces. For example, the *GSTM1* deletion and multiple *NAT* (N-acetyltransferase) variants were often considered together as the leading candidates for explaining the genetic basis of bladder cancer susceptibility (García-Closas *et al*. 2005). *NAT2* has been reported to be evolving under balancing selection (Mortensen *et al*. 2011). In addition, variations in other xenobiotic metabolism genes, such as members of the *SLC* (solute carrier) gene family, have also been shown to be evolving adaptively (Sabeti *et al*. 2007). Some of these variations involve gene deletions. For example, the deletion of *UGT2B17* (uridine diphosphoglucuronosyl-transferase) has been shown to have increased in frequency in East Asian populations under adaptive forces (Xue *et al*. 2008). Overall, functional variation affecting metabolizing genes may be maintained adaptively in the human populations due to varying environmental pressures.

### Implications to evolutionary medicine

As mentioned above, the variation in the *GSTM* locus has been the subject of more than 500 studies within the context of multiple diseases (Parl 2005). However, given that majority of these studies are correlations, they lack mechanistic or evolutionary insights as to why the genetic variation in this locus is relevant. The emerging picture is that of a gene family (*GSTM1-5*) that broadly metabolizes multiple carcinogenic substances, for example, 4-nitroquinoline-1-oxide (NQO) (Hayes *et al*. 2005). On top of that functional layer, it has been shown that this gene family is riddled with common genetic variation, including the unusually common deletion of the *GSTM1*. These genetic variants, as expected, were associated with multiple cancers (Parl 2005). The mechanistic explanation would be that the reduced the *GSTM* function leads to higher susceptibility to carcinogenic substance exposure (Hayes *et al*. 2005).

However, there seems to be some functional redundancy among the *GSTM* gene family members. As a consequence, the association between single variants to traits may not completely capture the biomedical impact of the variation in this locus as a whole. For example, the *GSTM2* may compensate some of the lost function due to the *GSTM1* deletion (Bhattacharjee *et al*. 2013). As such, the overall functional impact of a variant depends on its genomic context. Moreover, a recent pathway-level analysis revealed that glutathione conjugation pathway, for which the *GSTM* genes are central, is a significant player in determining breast cancer susceptibility (Wang *et al*. 2017). The implication being that rather than single variants, the combined effect of multiple variants affecting the function of the genes in this pathway eventually contributes to the overall disease susceptibility. This is not a new concept (Jin *et al*. 2014). But, we are now in a position to quantitatively address this issue, especially using genealogical approaches as we described for the *GSTM1* here.

From an evolutionary medicine point of view, it is important to highlight two interrelated features of the *GSTM* locus. First, even when a single haplotype affects susceptibility to disease, this may be due to pleiotropic effects. Indeed, we described multiple putative functional effects of the *Tanuki* haplogroup in East Asian populations, including the loss of function due to the *GSTM1* deletion and the decrease of the *GSTM5* expression. This result exemplifies the benefits of a haplotype-level understanding of the genetic variation. We argue based on our results and those of others (Liu *et al*. 2005; Claussnitzer *et al*. 2015) that the co-occurrence of multiple functionally and biomedically relevant variants in particular haplotypes should be treated as the norm, rather than the exception. As exemplified in this study, functional analysis of haplotypes that are under selection may provide crucial targets for future mechanistic studies.

Second, our results add to the growing list of population-specific haplotypes that may contribute to disease susceptibility, further underlying the importance of conducting genetic epidemiology studies in ancestrally diverse populations (Rosenberg *et al*. 2010; Wojcik *et al*. 2017). This is especially pertinent to the *GSTM* locus, given that environmental toxins and carcinogens, which may vary from one population to the other, are the primary target for this gene family. Such loci are also targets for local selection in humans, may be best exemplified by the recent study on Arsenic adaptation in Argentinian Andes population (Schlebusch *et al*. 2015).

## CONCLUSION

In this study, we report a particular haplogroup (*Tanuki* haplogroup) carrying the deletion allele that has likely evolved under non-neutral conditions and reached a high allele frequency in East Asian populations. This haplogroup has a broad regulatory effect on the metabolizing *GSTM* gene family. Overall, our study adds to the emerging notion that complex loci involving gene conversion events and structural variants may contribute to adaptive and biomedically relevant phenotypic variation (Boettger *et al*. 2016; Sekar *et al*. 2016).

## Acknowledgements

This study is supported by MS’s fund from Astellas Foundation for Research on Metabolic Disorders, Graduate Program for Leaders in Life Innovation, The University of Tokyo Life Innovation Leading Graduate School from MEXT, Japan and Grant-in-Aid for Japan Society for the Promotion of Science (JSPS) Fellows Grant Number 264456. This work is also supported by OG’s start-up funds from University at Buffalo Research Foundation.

We would like to thank Skyler Resendez for careful reading of the earlier versions of this manuscript. We would like to thank Takafumi Ishida, Jun Ohashi, Naoko Fujito and Mehmet Somel for discussions with regards to population genetic analyses. This study constitutes a part of the doctoral thesis of Marie Saitou submitted to The University of Tokyo, Japan.

